# Synergies analysis produces consistent results between motion analysis laboratories

**DOI:** 10.1101/2020.10.21.349126

**Authors:** Bruce A. MacWilliams, Mark L. McMulkin, Adam Rozumalski, Michael H. Schwartz

**Author notes:** Corresponding author, Motion Analysis Center, Shriners Hospitals for Children^®^, 1275 Fairfax Rd., Salt Lake City, UT 84103-4399.

## Abstract

**Aim:** The dynamic motor control index during walking (*walk-DMC*) has been shown to be related to patient outcomes and there has been an increasing interest from motion analysis centers regarding using this metric in their own practice. However, the methods for computing the index reported in the literature are not consistent. Here we propose a standardized method and investigate if this leads to results that are consistent between laboratories.

**Method:** Comparisons of typically developing controls are made between three independent motion analysis centers. Comparisons are also made between the proposed and previously published methods. A program script to compute the *walk-DMC* was used for this study and is made freely available with this manuscript.

**Results:** Using this script, results are highly consistent between three participating centers. The currently proposed method results in a wider distribution of *walk-DMC* values than those previously reported.

**Interpretation:** Using consistent processing methods, synergy measures are equivalent between centers. The major differences between current and published data are attributed to the use of concatenation of several walking trials.

---

The dynamic motor control index during walking (*walk-DMC*) measures the complexity of the lower extremity muscle activation sequence (1). This index is derived from a synergy analysis of the electromyographic (EMG) activity of muscles in the lower extremities recorded during walking. The variance accounted for by one synergy (*VAF_1_*) is computed, then scaled into a transformed *z*-score using normative values

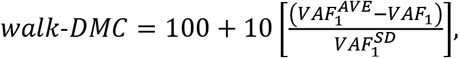

where scaling parameters *VAF_1_^AVE^* and *VAF_1_^SD^* are the means and standard deviations of *VAF_1_* from a typically developing population.

Because *VAF_1_* is a measure of the complexity of the signals explained by one synergy, a larger value of *VAF_1_* implies a less complex set of signals. It has been shown that *walk-DMC* can predict outcomes (2) and is repeatable within a center (3). It is currently unknown if *walk-DMC* values are comparable between centers using their own normative datasets to establish the *VAF_1_A^VE^* and *VAF_1_^SD^*. For centers implementing *walk-DMC* testing, it is important to establish the expected values and determine if center specific assessment of normative EMG is required, or if generic *VAF_1_* scaling values can be used. This compatibility is an essential component necessary to conduct multi-center research and compare individual center results to published values.

The methods used for computing *walk-DMC* have evolved over time, and currently there are several approaches in widespread use. For example, authors have previously used either four or five muscles per limb (1,2). Also, different signal processing techniques have been employed (3). The resulting *VAF_1_* and *walk-DMC* values may also be dependent on the number of strides analyzed, and whether these strides are averaged or concatenated.

This study addresses three main goals related to computing *walk-DMC*. The first goal is to establish how consistent *walk-DMC* values are between centers using their own normative data sets. This includes variability which arises from differences in the populations sampled, staff experience, protocol and equipment, and laboratory configurations and sizes. The first specific hypothesis tested is that *VAF_1_*^AVE^ and *VAF_1_^SD^* determined from locally collected typically developing populations will not be significantly different between multiple laboratories. The second goal is to determine if using a five muscle set of lower limb EMG data results in differences in *walk-DMC* compared to a four muscle set, which has been previously used. The second specific hypothesis tested is that *VAF_1_* using 5 muscle groups will be significantly lower than *VAF_1_* determined using 4 muscle groups, as an additional signal will add significant unexplained variance to the dataset. Finally, the third goal is to compare the *VAF_1_^AVE^* and *VAF_1_^SD^* scaling values determined from the current signal processing protocol to those used in two previous studies (1,2). The third and final hypothesis is that scaling factors determined by the proposed current data processing techniques will result in appreciable changes in *walk-DMC* values.

## Methods

Three independent motion analysis centers contributed data for analysis in this retrospective assessment (Shriners Hospitals for Children – Salt Lake City, Shriners Hospitals for Children – Spokane, Gillette Children’s Specialty Healthcare). Each center contributed a dataset derived from typically developing children between the ages of 4-21 collected at their own institution using their own equipment, staff, and protocols. All subjects were screened for recent orthopedic or neurological concerns. All protocols included placement of EMG preamplifiers bilaterally on the rectus femoris, vastus lateralis, medial hamstrings, anterior tibialis, and medial gastrocnemius/soleus. Similar equipment was used between laboratories (MA-300 or MA-400, Motion Lab Systems, Baton Rouge, LA). While no single formal standard protocol was employed in all laboratories, preamplifiers were placed using common general guidelines (4,5). All individuals walked at a self-selected speed along a ~10 *m* walkway. Five trials were used for each individual. The first and last 10% of each trial was trimmed to eliminate potential transient effects of acceleration and deceleration. Data was collected at 1000 *Hz* or 1080 *Hz*. A single software script (compatible with Matlab version 2016b and newer, with no required toolboxes) was used in each laboratory to process the typically developing data sets (Matlab, Natick, MA). This script, which assumes input from appropriately formatted C3D files (C3D.org), is included with this report. For motion analysis systems that do not produce C3D files, the file input structure of the script can be modified. The pertinent sequential signal processing steps of the raw data for each included file are:

1. **High pass filter**: Apply 4^th^ order zero lag Butterworth high pass (HP) 35 *Hz* filter.
2. **60 *Hz* noise assessment and filter**: Calculate the power in the bandwidth between 59-61 *Hz* in each channel. If this bandwidth power is > 10% of the entire signal power, apply 4^th^ order zero lag Butterworth notch filters for both the 59-61 *Hz* range and the 119-121 *Hz* range.
3. **Demean**: Adjust signals to give a zero mean value.
4. **Rectify**: Apply full wave rectification to the signals.
5. **Low pass filter**: Apply 4^th^ order zero lag Butterworth low pass (LP) 10 *Hz* filter.
6. **Trim**: Truncate first and last 10% of signal to remove filter end effects and periods of transient gait.
7. **Normalize**: Divide each channel by the maximum value of that channel. Sparsely occurring negative values are set to zero.
8. **Concatenate**: Splice together all included trials to create a single array encompassing many strides.

Note that filter coefficients are computed based on specified cutoff values and data frequency, which is automatically read from the data file. The *VAF_1_* from these resulting signals is then computed using previously published techniques.

Data from both limbs of each individual were included in the analysis. The *VAF_1_* distributions between laboratories were compared using ANOVA, and distribution variance was compared using Bartlett’s test. Additionally, *VAF_1_* distributions between the four and five muscle calculations were compared; mean values were compared using Student t-tests, and variances were compared using Bartlett’s test. An alpha level of 0.05 was used for all statistical hypothesis testing, and all statistical analyses were performed in Matlab (version 2020a, Mathworks, Natick, MA). An analysis of the current processing technique was compared to previously published techniques using theoretical minimum and maximum values, and *walk-DMC* values computed from 350 subjects with cerebral palsy [2]. Scaling values reported in two prior publications were used here for comparison, one determined from a five muscle data set and a single stride (1), and one from a four muscle data set using the average of between 3 and 6 walking trials (2).

## Results

### Consistency between centers

All three participating centers utilized pediatric populations to establish normative EMG data sets, and there were no marked differences between the demographics (Table 1). There were no significant differences between the laboratories in either *VAF_1_^AVE^* or *VAF_1_^SD^* (Table 2). This was true using either the four muscle data set or the five muscle data set.

Consistency Between 4 and 5 Muscles

**Table 1.** Demographic data of the typically developing individuals used at each laboratory to determine *VAF_1_^AVE^* and *VAF_1_^SD^. N* = number of participants. Speed is determined through the instrumented gait data of the trials used in this analysis. Continuous variables are given as mean ± standard deviation.

| Center | N | Gender (F/M) | Age (Y) | Height (cm) | Weight (kg) | BMI | Speed (m/s) |
| --- | --- | --- | --- | --- | --- | --- | --- |
| Center1 | 39 | 22/17 | 13.6 ± 4.4 | 152.8 ± 17.3 | 51.3 ± 18.5 | 21.2 ± 4.2 | 1.2 ± 0.1 |
| Center2 | 29 | 19/10 | 9.0 ± 2.7 | 133.6 ± 15.6 | 31.1 ± 11.0 | 16.9 ± 2.5 | 1.2 ± 0.1 |
| Center3 | 24 | 9/15 | 11.7 ± 3.6 | 150.0 ± 21.7 | 44.5 ± 19.2 | 18.8 ± 3.6 | 1.3 ± 0.2 |

**Table 2.** Comparison between laboratory normative control sets. *N* = number of limbs. The first three rows show individual laboratory results, the 4^th^ row is the combined data set. The 5^th^ row reports p value of *VAF_1_^AVE^* from ANOVA between the three labs, and p value of *VAF_1_^SD^* from Bartlett’s test of variance. The final two columns compare the 5 muscle data set to the 4 muscle data set using Student t-tests for *VAF_1_^AVE^* and Bartlett’s test of variance for *VAF_1_^SD^*.

| Center | 5 Muscles |  |  | 4 Muscles |  | 5 vs. 4 Muscles |  |
| --- | --- | --- | --- | --- | --- | --- | --- |
| | N | $VA_{F_1}^{AVE}$ | $VA_{F_1}^{SD}$ | $VA_{F_1}^{AVE}$ | $VA_{F_1}^{SD}$ | p $VA_{F_1}^{AVE}$ | p $VA_{F_1}^{SD}$ |
| Center1 | 78 | 0.6683 | 0.0451 | 0.6639 | 0.0460 | 0.5481 | 0.8680 |
| Center2 | 58 | 0.6761 | 0.0404 | 0.6675 | 0.0429 | 0.2678 | 0.6576 |
| Center3 | 48 | 0.6648 | 0.0409 | 0.6812 | 0.0447 | 0.1456 | 0.5460 |
| Combined | 184 | 0.6708 | 0.0425 | 0.6696 | 0.0450 | 0.7872 | 0.4425 |
| p Between Labs |  | 0.5200 | 0.6126 | 0.1009 | 0.8545 |  |  |

### 4 versus 5 muscles

There were no differences in *VAF_1_^AVE^* between five muscles per limb and four muscles per limb in any laboratory, or between the combined data set. There were also no differences in *VAF_1_^SD^*, as determined by Bartett’s test.

### Comparison to previous techniques

Using the methods proposed here results in greater ranges of *walk-DMC* theoretical minimums and maximums compared to previous single stride and trial averaging approaches (Table 3). This effect is additionally reflected in the *walk-DMC* variance of 350 subjects with cerebral palsy compared to the same set of data compiled using either a single stride or the mean of several trials (Figure 1). Both the means and variances of the distributions are significantly different (p<0.0001). For means, the single stride approach led to a significantly lower value than the other methods. For variances, the concatenated stride approach leads to a wider distribution of *VAF_1_* values than the other approaches. In this dataset using the concatenated approach, mean *walk-DMC* values for GFMCS I, II, III subjects were 92.8, 72.6, 67.2 respectively and ranged from 46.0 to 121.3.

**Table 3.**
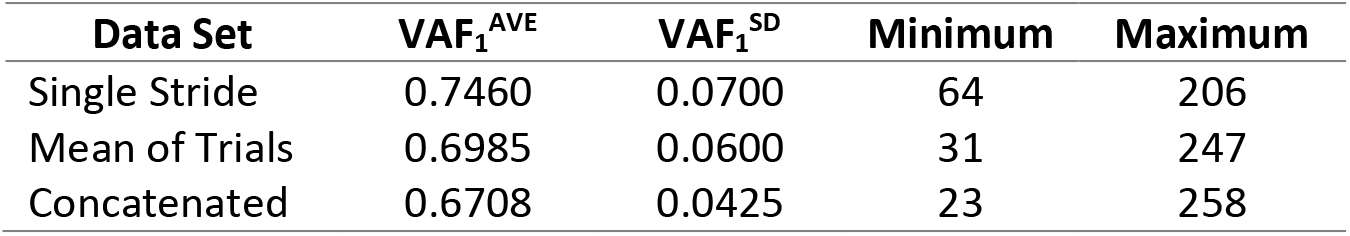
Theoretical minimum (setting *VAF_1_* = 1) and maximum (setting *VAF_1_* = 0) values of *walk-DMC* as determined by scaling values *VAF_1_^AVE^* and *VAF_1_^SD^*. The concatenated values used here are the combined values of the participating labs (Table 2, row 4).

**Figure 1.**
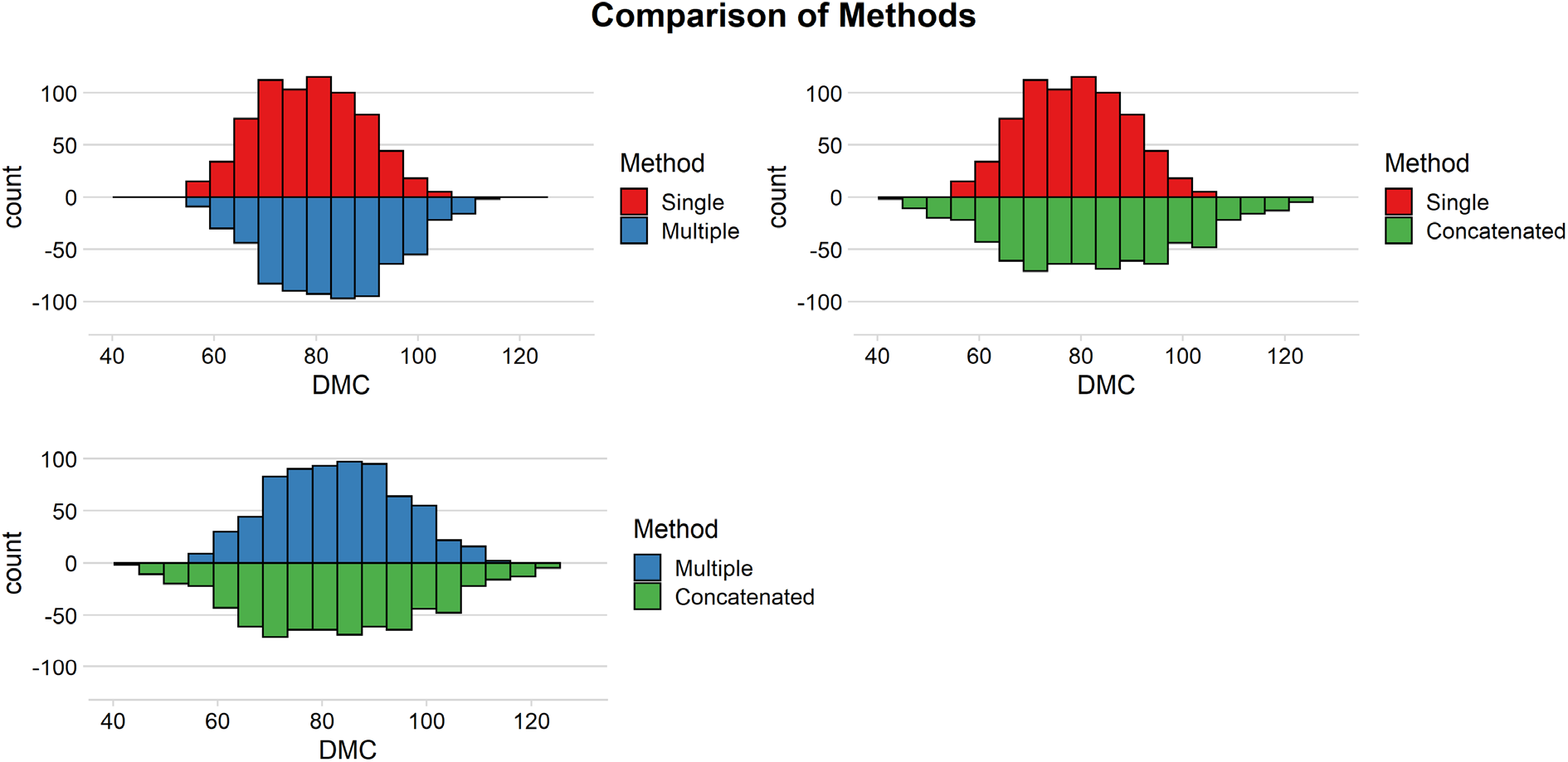
Comparisons of distributions of *walk-DMC* of 350 children with cerebral palsy from different processing techniques used to compute the *walk-DMC*. Red distribution is from previous work using a single stride for each typically developing individual to determine *VAF_1_^AVE^* and *VAF_1_^SD^*. Blue distribution results from using a mean of several trials. The broader green distribution utilizes the concatenation of several trials as reported in the current work.

## Discussion

The purpose of this study was to assess the repeatability of the *walk-DMC* between labs, introduce recommended data collection and processing techniques, and provide free open-source code for calculation of *VAF_1_* using the recommended approach. The means and variances of *VAF_1_* distributions did not differ between three centers that independently collected data. This supports the first hypothesis that, with a consistent processing approach, differences between underlying typically developing datasets are not important to synergy decomposition. In addition, whether four or five muscles were used did not matter. This refutes the second hypothesis, that the addition of the vastus lateralis would add additional unexplained variance. Finally, current processing technique changes, particularly the use of concatenated trials, led to a wider distribution of *Walk-DMC* values compared to previous reports, supporting the third hypothesis that signal processing techniques need to be consistently implemented to obtain consistent *walk-DMC* distributions.

Motivated by concerns about inter-laboratory differences in data processing, Shuman *et al*. reported that the *z*-score implementation inherent in the *walk-DMC* calculations reduced sensitivity to low pass filter choices compared to raw *VAF_1_* values (3). In this study, we demonstrate that when using identical signal processing techniques, *VAF_1_* is not significantly different between laboratories. Both the means and the variances of the *VAF_1_* distributions were equivalent between the three centers, such that *VAF_1_^AVE^* and *VAF_1_^SD^* and calculated *walk-DMC* values are also equivalent.

It is important to determine how the choice of different sets of EMG channels may affect *walk-DMC,*since standards may vary between centers. In this work we compared a four muscle set used in previous work to a five muscle set not previously used for *walk-DMC* calculations, but which is the current standard for clinical analyses in all three of the participating laboratories. If the additional EMG channel captures additional information regarding neurologic control, then it will decrease *VAF_1_*. The value of *VAF_1_^AVE^* did not differ between four and five muscle sets (<2.5%, not statistically different). Thus, one finding from this study is that the four muscle set is adequate, and indeed equivalent to the five muscle set.

For the CP population studied here, *walk-DMC* values stratified by GMFCS levels I-III were 92.8, 72.6, 67.2 on average respectively and ranged from approximately 40-125 (Figure 1). In a previous report, Schwartz *et al*. demonstrated that pre-operative *walk-DMC* values were associated with treatment outcomes in children with cerebral palsy (2). Plots indicate the range of *walk-DMC* for the 473 GMFCS IIII individuals in the outcome study ranged between 60 and 100 using *VAF_1_^AVE^* and *VAF_1_^SD^* derived from data collected from four muscles and averages of multiple trials, each of which contained several strides. Similarly, Steele *et al*. reported *walk-DMC* values stratified by GMFCS level in 633 individuals with cerebral palsy (1). Stratified means of GMFCS I, II, and III were 92.4, 85, and 82, respectively. These were computed from *VAF_1_^AVE^* and *VAF_1_^SD^* derived from a five muscle set that did not include a vastus muscle, and used just one randomly selected gait cycle. In the current work, a different five muscle set is used and five trials are concatenated, yielding a minimum of 10 strides per individual. The inclusion of many more strides for analysis reduces normative *VAF_1_* variance (*VAF_1_^SD^*), and therefore magnifies the differences between *VAF_1_^CP^* and *VAF_1_^AVE^*. Although study populations were different, the effect of basing *walk-DMC* calculations on a single stride, versus several strides, versus many strides is clearly reflected in the different ranges and GMFCS stratified means found. The comparisons of theoretical minimum and maximum values for the three sets reinforce this concept. For example, the theoretical *walk-DMC* minimum of single stride values (64) is markedly higher than the minimum of combined data set using five concatenated trials (23), effectively limiting the range of possible clinical values. Using the proposed processing techniques widens the expected range of *walk-DMC* values (Table 3). Alternatively stated, *walk-DMC* values exhibit a wider distribution when using the concatenated approach (Figure 1).

There are several recognized limitations to this study. Though data suggest that concatenated analysis leads to a wider distribution of *walk-DMC* values than previous methods, we have not assessed how many trials or strides are needed to reach a stable value. Also, this work proposes, but does not explore the effects of filter values. Previous work reported that *VAF_1_* and *walk-DMC* were most affected by LP filter choice compared to other signal processing choices. Increased LP cutoff frequencies may artificially inflate the apparent complexity of the synergy, while if an LP cutoff is set too low, actual complexity in the signals may be lost. In this work, we chose to use a 10 *Hz* filter based on Shuman *et al*. (3). There are many factors in the clinical protocol which may lead to potential differences in *VAF_1_* distributions and *walk-DMC* values. While this study found no differences between the three participating independent centers it should be noted that these three centers have similarly experienced personnel, equipment, and a history of informal collaboration which may have influenced these findings.

This study has shown that *VAF_1_^AVE^* and *VAF_1_^SD^* for typically developing individuals were not significantly different between centers. With consistent implementation of the same muscles tested and signal processing techniques, *VAF_1_* (and therefore, *walk-DMC*) values can be compared between centers. Additional centers implementing *walk-DMC* analysis from their own typically developing EMG data should expect similar results. Alternatively, a center not collecting their own typically developing EMG data should be able to implement *walk-DMC* calculations for clients with cerebral palsy by using the *VAF_1_^AVE^* and *VAF_1_^SD^*, testing the same muscles, and exact processing protocol from this paper. Centers are still encouraged to collect and utilize locally collected typically developing data to establish these values as this is recognized as the best practice and there may be additional factors including staff experience, placement, and equipment that may lead to potential differences to the values determined here. This is particularly true in light of the similarities in experience, protocols, and equipment between the three centers participating in the current study. To facilitate this effort, and to give centers the ability to calculate their own scaling factors as well as subject values, we have provided a Matlab script with this article as well as instructions on its use. The script allows flexible output and simple implementation but also includes some features for more advanced analysis such as exploring additional synergy levels and enabling users to visually interpret the effects of each filter processing step. While the script is built to accept C3D files, it could be altered by any experienced Matlab user to input other file types.

## Notes

### Competing Interest Statement

The authors have declared no competing interest.

## References

1. Steele KM, Rozumalski A, Schwartz MH. Muscle synergies and complexity of neuromuscular control during gait in cerebral palsy. Developmental Medicine & Child Neurology. 2015 Dec 1;57(12):1176–82.

2. Schwartz MH, Rozumalski A, Steele KM. Dynamic motor control is associated with treatment outcomes for children with cerebral palsy. Dev Med Child Neurol. 2016 Nov;58(11):1139–45.

3. Shuman BR, Schwartz MH, Steele KM. Electromyography Data Processing Impacts Muscle Synergies during Gait for Unimpaired Children and Children with Cerebral Palsy. Front Comput Neurosci. 2017;11:50.

4. Perotto AO. Anatomical Guide for the Electromyographer: The Limbs and Trunk. 4th ed. Springfield, IL: Charles C. Thomas; 2004. 345 p.

5. Hermens HJ, Freriks B, Merletti R, Stegeman D, Blok J, Rau G, et al. Surface ElectroMyoGraphy for the Non-Invasive Assessment of Muscles [Internet]. 1999. Available from: www.seniam.org

